# Fast-Bonito: A Faster Basecaller for Nanopore Sequencing

**DOI:** 10.1101/2020.10.08.318535

**Authors:** Zhimeng Xu, Yuting Mai, Denghui Liu, Wenjun He, Xinyuan Lin, Chi Xu, Lei Zhang, Xin Meng, Joseph Mafofo, Walid Abbas Zaher, Yi Li, Nan Qiao

## Abstract

Oxford Nanopore Technologies (ONT) is a promising sequencing technology that could generate relatively longer sequencing reads compared to the next generation sequencing (NGS) technology. The base calling process is very important for TGS. It translates the original electrical signals from the sequencer to the nucleotide sequence. By doing that, the base calling could significantly influence the accuracy of downstream analysis. Bonito is a recently developed basecaller based on deep neuron network, the neuron network architecture of which is composed of a single convolutional layer followed by three stacked bidirectional GRU layers. Although Bonito achieved the state-of-the-art accuracy, its speed is so slow that it is not likely to be used in production. We therefore implement Fast-Bonito, which introduces systematic optimization to speed up Bonito. Fast-Bonito archives 53.8% faster than the original version on NVIDIA V100 and could be further speed up by HUAWEI Ascend 910 NPU, achieving 565% faster than the original version. The accuracy of Fast-Bonito is also slightly higher than the original Bonito.

## Introduction

In the last several decades, genomic sequencing technologies have been evolved from Sanger sequencing^1^ to massively parallel next generation sequencing (NGS)^2^, and long-read third-generation sequencing (TGS)^3^. Compared to NGS, TGS could produce longer reads, which make it a better choice to study complex genome and variants. Several different TGS technologies have been developed, such as PacBio and Oxford Nanopore Technology (ONT) platform^3^. Nanopore sequencing powered by ONT could generate very long reads by saving the electrical resistance signals of the single strand DNA that passing through the protein nanopore^4^. It shows great advantage on sequencing long reads and detecting complex genome structure variation^4,5^, but the high sequencing error rates slowed down its industry use in many areas.

A very important step that introduces the high error rates is base calling. This process is used to translate the raw electrical signal into nucleotide sequence. Although a number of machine learning approaches have been used for base calling^6–12^, it is still quite challenging to obtain a fast and accurate basecaller. The electrical signals are determined by multiple nucleotides residing in the nanopores, but the noises, such as the emergence of DNA methylation will make the signals very complex^12^ to decode.

Many approaches have been developed for the base calling task. For example, ONT released Albacore, Guppy, Scappie, Flappie (https://github.com/nanoporetech/flappie) and Bonito (https://github.com/nanoporetech/bonito); research communities also contribute to a lot of tools, such as Nanocall^10^, DeepNano^9^, Chrion^8^ and causalcall^7^. In recent years, deep neuron network has been used more and more in tools for this task, including convolutional neural network (CNN), recurrent neural network (RNN) and connectionist temporal classification (CTC) decoder^7^. These tools vary a lot in speed or accuracy. And currently Guppy is an order of magnitude^12^ faster than all the others, also with a relatively high accuracy.

Recently, a new algorithm, Bonito, has been developed and achieved state-of-the-art accuracy, representing a significant improvement of over 1% comparing to Guppy (https://nanoporetech.com/about-us/news/new-research-algorithms-yield-accuracy-gains-nanopore-sequencing). The speed of Bonito, however, is very slow, which limits its application in practice. Therefore, we developed Fast-Bonito, which introduces systematic approaches to optimize the architecture of Bonito, and achieves 565% faster sequencing speed than the original version.

## Results

### Search for an Optimized Neuron Network Backbone

The neural network backbone of Bonito is inspired by QuartzNet^13^, which is originally developed for speech recognition. The backbone of Bonito consists of several TCSConv-BN-ReLU modules, each contains a depth-wise separable convolution. It reduces the number of parameters of the model dramatically. However, hardware and inference engine library cannot support such convolution operator well, which makes its running time slower than expectation.

To speed up Bonito, we use neural architecture search (NAS) to search for a more efficient neuron network for the same task. Previous NAS frameworks only focus on searching for a higher performance module, which might result in a backbone that satisfies the accuracy, but takes longer time for inference (higher latency). To balance both the accuracy and inference latency, a multi-objective and adaptive NAS framework (so called MnasNet^14^) was used in our work. The original separable convolution module was replaced by Bottleneck Convolution module from ResNet50^15^, because the latter is more friendly supported by current inference engine library. The NAS search focused on the architecture of five middle blocks. The search space of each block was defined as follows. The number of modules of each block is selected from 1 to 9.

1. The number of channels of block is searched from 32, 64, 128, 256, 378, 512.
2. The kernel size of Bottleneck Conv operator is searched from 3, 5, 7, 9, 11, 17, 29, 31, 47, 53, 69, 73, 83, 91, 107, 115, 123, 129.

The best backbone from NAS search is named Fast-Bonito, the overall architecture is shown in Fig. 1, including a sequential of convolution module layers to calculate the probability of nucleotide, and a CTC decoder to translate the probability into nucleotide sequence. Considering that the electrical signals are diverse in length, we also performed segments-split, referring to Bonito. The input signals are cut into 6,000 segments with an overlap of 300 before fed into the convolutional architecture, with corresponding output segments with length of 6,000. Both ends of segments are removed by a length of 150 except the first and last one. The back-end and leading-end of the first and last segments are removed, respectively. The remaining parts of segments are concatenated together and fed into the CTC decoder to produce final nucleotide sequencing.

**Figure 1.**
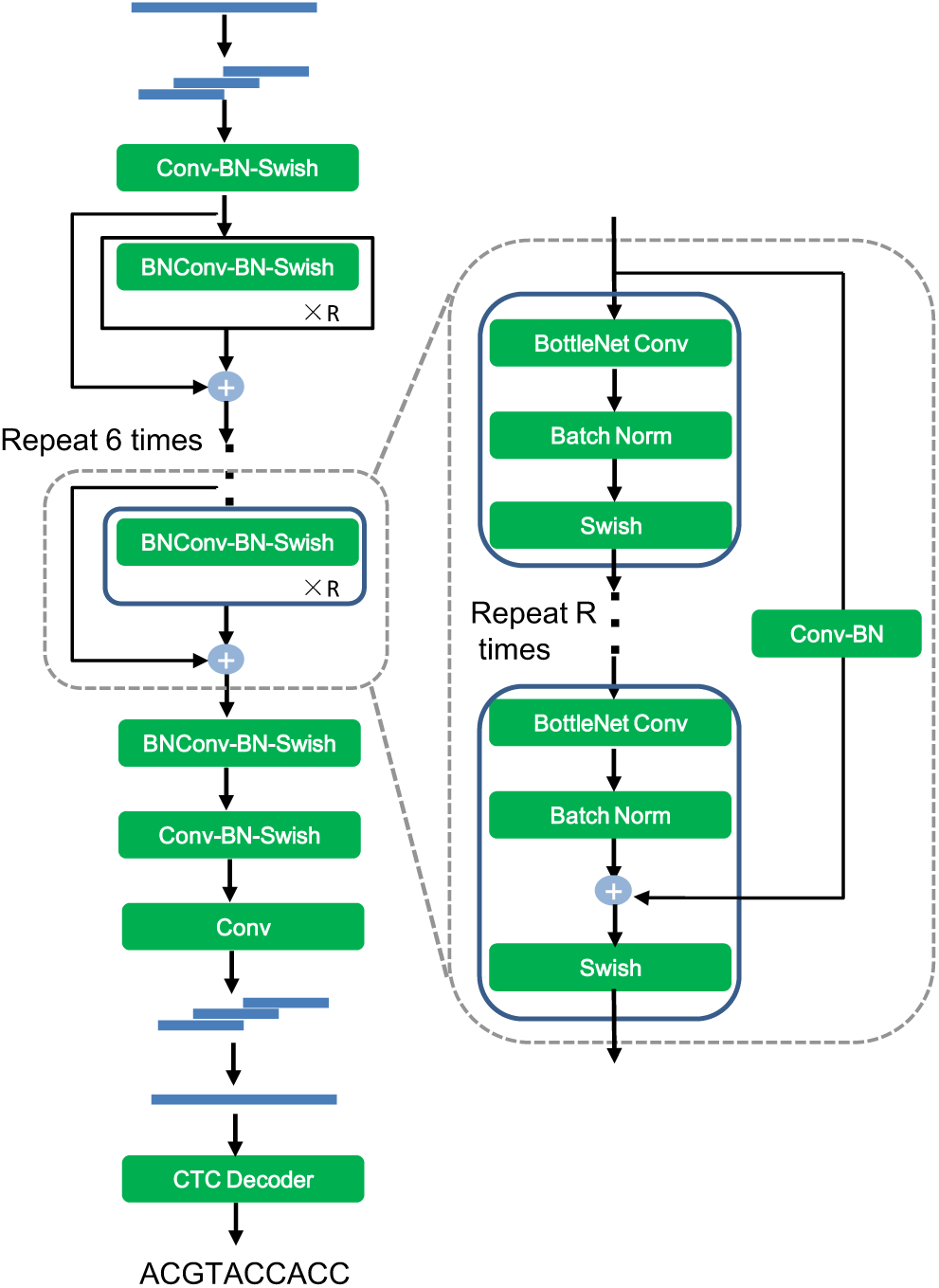
Backbone of Fast-Bonito. Fast-Bonito replaced the Separable Convolution module in Bonito by Bottleneck Convolution module for acceleration. The architecture of Fast-Bonito comprises a convolutional architecture, a Convolution module layer in the last layer that calculates the probability of nucleotide, and a CTC decoder to translate the output of network into nucleotide sequence.

### Train New Backbone from Scratch

After the best backbone of Fast-Bonito is decided, we began to train it from scratch using the optimization tricks below.

1. Data Augmentation is a popular method to improve the model performance^16–18^. The common augmentation approaches, such as cutout^19^, rotate, flip^20^, shearX^21^, are not very suitable for 1-dimensional DNA sequence. We therefore used SpecAugment^18^, a data augmentation method for speech recognition, which is similar to DNA sequence decoder.
2. Label Smoothing^22,23^ is a method to prevent the network becoming over-confident and has been utilized in many state-of-the-art models. Here we introduced this method to prevent over-fitting.
3. Knowledge Distillation^24,25^ is a method to migrate the knowledge from the well-trained “teacher” model to the “student” model, thus to improve the performance of “student” model. Here we take the pre-trained model of the original Bonito as “teacher” and our own model as “student”.

### Model performance comparison

Our primary benchmark dataset that is used to estimate the speed and accuracy of Fast-Bonito is obtained from the dataset provided by Bonito. This dataset contains 100000 reads for evaluation. On NVIDIA V100 GPU. The speed of the original version of Bonito is 1,400,000 bp/s (Fig. 2A). Fast-Bonito could achieve 1,840,000 bp/s in speed, which is 53.8% faster than the original Bonito.

**Figure 2.**
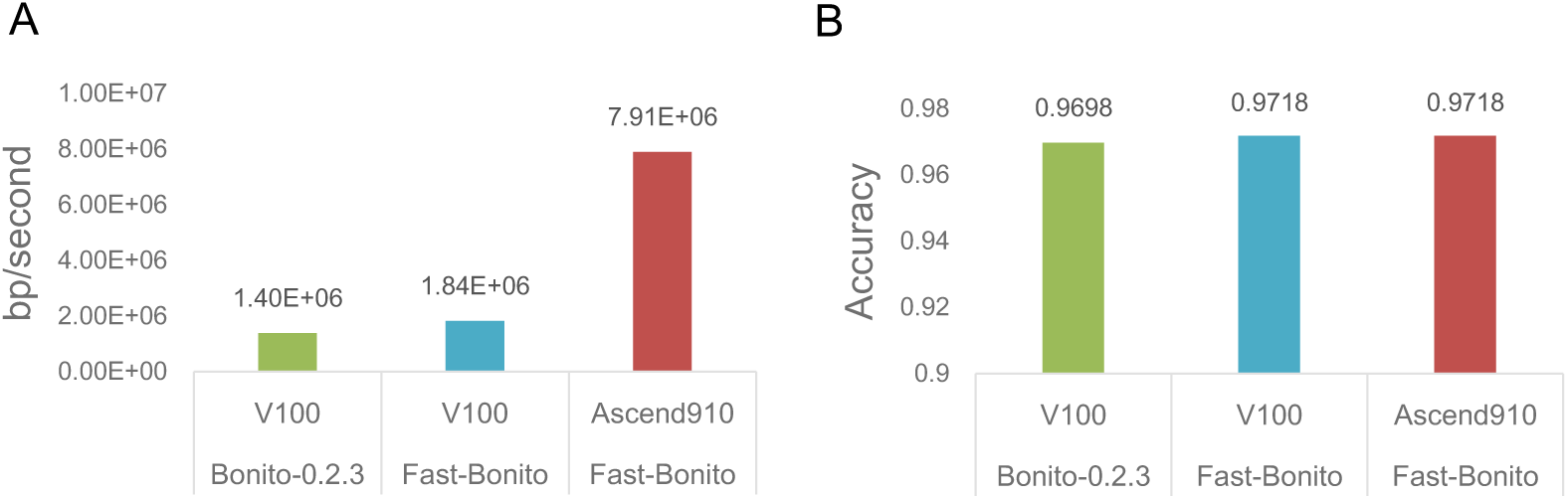
Performance of FastBonito. A). On NVIDIA V100, Fast-Bonito 0.54 times faster than the original version. While on Asscend 910 NPU, Fast-Bonito is 4.65 times faster. B). The accuracy of basecalling of Fast-Bonito is slightly higher than the Original-Bonito (by 0.2%).

As Neuron Process Unite (NPU) is more and more popular for AI computing, we also tested Fast-Bonito on HUAWEI Ascend 910 Chip, an NPU from HUAWEI. The results show that on HUAWEI Ascend 910 Chip, Fast-Bonito could achieve 7,910,000 bp/s in speed, which is 565% faster than the original version (Fig. 2A).

Fast-Bonito could achieve a slightly better median accuracy, which is 97.18% on NVIDIA V100 and HUAWEI Ascend 910 (Fig. 2B), compared to 96.98% of the original Bonito.

## Discussion

Base calling is the key step for the nanopore sequencing analysis workflow. Fast and accurate base calling is still a challenging problem. Although Bonito achieves the state-of-the-art accuracy, the speed of Bonito limits its application into the production. Base calling tools require huge computational resources, especially GPU resources. Our study demonstrates that AI processor could be used to accelerate base calling, and could achieve 4.3 times faster than on NVIDIA V100 GPU, indicating the promising use of AI processor on genomic study.

The Bonito version in this study is 0.2.3, Fast-Bonito is also developed with the same training and validation dataset. Bonito is still an active project, and continuously releases new features. We will also continuously update Fast-Bonito with the new features in the future.

## Data and Software Statement

Training and validation data is downloaded from GitHub of Bonito. Fast-Bonito is made public under https://github.com/EIHealth-Lab/fast-bonito.

### Test environment

The test environment of V100 is under the NVIDIA Tesla, 16 GB GPU with 8 CPUs. And the environment of Ascend 910 is under 16 GB*8 NPUs with 192 CPUs, while only 1 NPU and 24 CPUs are specified for current task.

### Metrics for Base calling Evaluation

To evaluate base calling, a pairwise sequence alignment library, parasail^26^, was used to calculate the accuracy between sequencing and referencing. “bonito evaluate dna_r9.4.1 –chunks 100000” was used to evaluate the performance for the original bonito model.

## Author Contributions

N.Q. and Y.L. designed and conceived the project. Y.M. performed the optimization of Bonito and model training. Z.X. and D.L. implemented the Fast-Bonito package and performed data experiment with the help from W.H, C.X., L.Z. and N.Q, D.L. wrote the manuscript. X.M., J.M. and W.A.Z. provided suggestions and revised the manuscript. All authors read and approved the final manuscript.

## Reference

1. Sanger, F. & Coulson, A. R. A rapid method for determining sequences in DNA by primed synthesis with DNA polymerase. J. Mol. Biol. 94, 441–448 (1975).

2. Behjati, S. & Tarpey, P. S. What is next generation sequencing? Arch. Dis. Child. - Educ. Pract. Ed. 98, 236–238 (2013).

3. Lee, H. et al. Third-generation sequencing and the future of genomics. http://biorxiv.org/lookup/doi/10.1101/048603 (2016) doi:10.1101/048603.

4. Mikheyev, A. S. & Tin, M. M. Y. A first look at the Oxford Nanopore MinION sequencer. Mol. Ecol. Resour. 14, 1097–1102 (2014).

5. Ho, S. S., Urban, A. E. & Mills, R. E. Structural variation in the sequencing era. Nat. Rev. Genet. 21, 171–189 (2020).

6. Huang, N., Nie, F., Ni, P., Luo, F. & Wang, J. An attention-based neural network basecaller for Oxford Nanopore sequencing data. in 2019 IEEE International Conference on Bioinformatics and Biomedicine (BIBM) 390–394 (IEEE, 2019). doi:10.1109/BIBM47256.2019.8983231.

7. Zeng, J. et al. Causalcall: Nanopore Basecalling Using a Temporal Convolutional Network. Front. Genet. 10, 1332 (2020).

8. Teng, H. et al. Chiron: translating nanopore raw signal directly into nucleotide sequence using deep learning. GigaScience 7, giy037 (2018).

9. Boža, V., Brejová, B. & Vinař, T. DeepNano: Deep recurrent neural networks for base calling in MinION nanopore reads. PLOS ONE 12, e0178751 (2017).

10. David, M., Dursi, L. J., Yao, D., Boutros, P. C. & Simpson, J. T. Nanocall: an open source basecaller for Oxford Nanopore sequencing data. Bioinformatics 33, 49–55 (2017).

11. Silvestre-Ryan, J. & Holmes, I. Pair consensus decoding improves accuracy of neural network basecallers for nanopore sequencing. http://biorxiv.org/lookup/doi/10.1101/2020.02.25.956771 (2020) doi:10.1101/2020.02.25.956771.

12. Wick, R. R., Judd, L. M. & Holt, K. E. Performance of neural network basecalling tools for Oxford Nanopore sequencing. Genome Biol. 20, 129 (2019).

13. Kriman, S. et al. QuartzNet: Deep Automatic Speech Recognition with 1D Time-Channel Separable Convolutions. ArXiv191010261 Eess (2019).

14. Tan, M. et al. MnasNet: Platform-Aware Neural Architecture Search for Mobile. in 2019 IEEE/CVF Conference on Computer Vision and Pattern Recognition (CVPR) 2815–2823 (IEEE, 2019). doi:10.1109/CVPR.2019.00293.

15. He, K., Zhang, X., Ren, S. & Sun, J. Deep Residual Learning for Image Recognition. ArXiv151203385 Cs (2015).

16. Cubuk, E. D., Zoph, B., Mane, D., Vasudevan, V. & Le, Q. V. AutoAugment: Learning Augmentation Strategies From Data. in 2019 IEEE/CVF Conference on Computer Vision and Pattern Recognition (CVPR) 113–123 (IEEE, 2019). doi:10.1109/CVPR.2019.00020.

17. Zoph, B. et al. Learning Data Augmentation Strategies for Object Detection. ArXiv190611172 Cs (2019).

18. Park, D. S. et al. SpecAugment: A Simple Data Augmentation Method for Automatic Speech Recognition. in Interspeech 2019 2613–2617 (ISCA, 2019). doi:10.21437/Interspeech.2019-2680.

19. DeVries, T. & Taylor, G. W. Improved Regularization of Convolutional Neural Networks with Cutout. ArXiv170804552 Cs (2017).

20. Shorten, C. & Khoshgoftaar, T. M. A survey on Image Data Augmentation for Deep Learning. J. Big Data 6, 60 (2019).

21. Hu, B., Lei, C., Wang, D., Zhang, S. & Chen, Z. A Preliminary Study on Data Augmentation of Deep Learning for Image Classification. ArXiv190611887 Cs Eess (2019).

22. Kim, S., Seltzer, M. L., Li, J. & Zhao, R. Improved training for online end-to-end speech recognition systems. ArXiv171102212 Cs (2018).

23. Szegedy, C., Vanhoucke, V., Ioffe, S., Shlens, J. & Wojna, Z. Rethinking the Inception Architecture for Computer Vision. in 2016 IEEE Conference on Computer Vision and Pattern Recognition (CVPR) 2818–2826 (IEEE, 2016). doi:10.1109/CVPR.2016.308.

24. Hinton, G., Vinyals, O. & Dean, J. Distilling the Knowledge in a Neural Network. ArXiv150302531 Cs Stat (2015).

25. Wei, L. et al. Circumventing Outliers of AutoAugment with Knowledge Distillation. ArXiv200311342 Cs (2020).

26. Daily, Jeff. Parasail: SIMD C library for global, semi-global, and local pairwise sequence alignments. BMC Bioinformatics, 17(1), 1–11. doi:10.1186/s12859-016-0930-z (2016).

